# Enterotypes of the human gut mycobiome

**DOI:** 10.1101/2022.12.13.520343

**Authors:** Senying Lai, Yan Yan, Yanni Pu, Shuchun Lin, Jian-Ge Qiu, Bing-Hua Jiang, Marisa Keller, Mingyu Wang, Peer Bork, Wei-Hua Chen, Yan Zheng, Xing-Ming Zhao

**Author notes:** Corresponding authors: Xing-Ming Zhao, Yan Zheng, Wei-Hua Chen, Peer Bork.

## Abstract

The fungal component of the human gut microbiome, also known as the mycobiome, plays a vital role in intestinal ecology and human health. Here, we identify and characterize four mycobiome enterotypes using ITS profiling of 3,363 samples from 16 cohorts across three continents, including 572 newly profiled samples from China. These enterotypes exhibit stability across populations and geographical locations and significant correlation with bacterial enterotypes. Particularly, we notice that fungal enterotypes have a strong age preference, where the enterotype dominated by *Candida* (i.e., fun_C_E enterotype) is enriched in the elderly population and confers an increased risk of multiple diseases associated with compromised intestinal barrier. In addition, bidirectional mediation analysis reveals that the fungi-contributed aerobic respiration pathway associated with fun_C_E enterotype might mediate the association between the compromised intestinal barrier and aging.

**Teaser:** As an integral part of the human gut microbiome, the fungi, which co-habit with prokaryotic microbiome in the gut, play important role in the intestinal ecology and human health. Yet, the overall structure of the human gut mycobiome and the inter-individual variation worldwide remain largely unclear. *Lai* et al. analyzed the fungal profiles of 3,363 samples from 16 cohorts across three continents, and identified four fungal enterotypes that exhibit stability across populations. They found that fungal enterotypes showed age preference, where a *Candida* dominated enterotype was enriched in the elderly population and confers an increased risk of multiple diseases and more severe compromised intestinal barrier. Furthermore, they determined one fungi-contributed aerobic respiration pathway could mediate the association between the compromised intestinal barrier and aging.

## Introduction

The human gut microbiome, which consists of multi-kingdom microbes of prokaryotes, viruses, protists and fungi, is essential to human health(*1*). Current research mainly focuses on the prokaryotic and viral components of the gut ecology(*2-4*). However, the complicated associations of other types of microorganisms, particularly fungi, with human health remain largely unknown. Although the fungal community, also known as mycobiome, comprises less than 1% of the entire human gut microbiome(*5*), they have been shown to be involved in disease pathogenesis and profoundly influence the host immune system(*6, 7*). For example, *Candida albicans* can cause infections in immunocompromised human hosts(*8*), and alterations of the gut mycobiome composition have been reported in multiple human diseases(*9, 10*). While fine-grained fungal taxonomic markers associated with certain phenotypes have been reported(*9, 11, 12*), the overall structure of the gut mycobiome and the inter-individual variation in fungal composition remain unclear.

Enterotypes, which have been proposed to summarize the human gut microbial characteristics, are effective in stratifying populations and providing a global overview of the inter-individual variations in gut microbial composition(*13, 14*). Multiple studies have consistently identified bacterial enterotypes, which were independent of the distribution of the hosts’ age, geography, and gender(*13-16*). Defined based on the prokaryotic compositional patterns, the enterotypes could enhance understanding of human health and facilitate intervention(*17*). As an integral part of the human gut multi-kingdom microbiome, the fungi share microhabitats with the prokaryotic microbiome in the gut through different types of interactions, such as mutualism, commensalism, and competition(*18*). Hence, they are important in shaping the bacterial community’s intestinal ecology. However, the landscape of the human gut mycobiome and whether fungal enterotype-like structures exist in the human gut are unclear.

In this study, we collected 3,363 fungal sequencing samples from 16 cohorts across Europe, North America, and Asia, including our 572 newly sequenced samples from China. Four fungal enterotypes were identified independent of populations and closely correlated with bacterial enterotypes. We noticed strong effects of host phenotypes (including age and diseases) on the fungal enterotypes. Notably, the *Candida* (fun_C_E) enterotype enriched in the elderly population showed a higher prevalence in patients of multiple diseases, even beyond the age influence, and was associated with a severe compromised intestinal barrier. Furthermore, a fun_C_E-enriched aerobic respiration pathway mediated the association between the compromised intestinal barrier and aging. Overall, our findings elucidated the highly structured nature of the gut mycobiome and its clinical relevance to human health.

## Results

### Landscape of human gut mycobiome composition and diversity

To characterize the human gut fungal diversity and composition, we collected internal transcribed spacer (ITS) sequencing data from 15 published projects (Supplementary Table 1)(*12, 19-28*). In addition, we recruited 572 Chinese participants (Chinese Gut Mycobiome cohort, or CHGM) aged from 17 to 89 years old and profiled their fecal mycobiome with ITS1 sequencing. In total, 3,363 fecal samples with ITS1- (960 samples) and ITS2- (2,403 samples) sequencing data from 16 cohorts covering three continents (Europe, North America, and Asia) were included in our study (Fig. 1a).

**Fig. 1.**
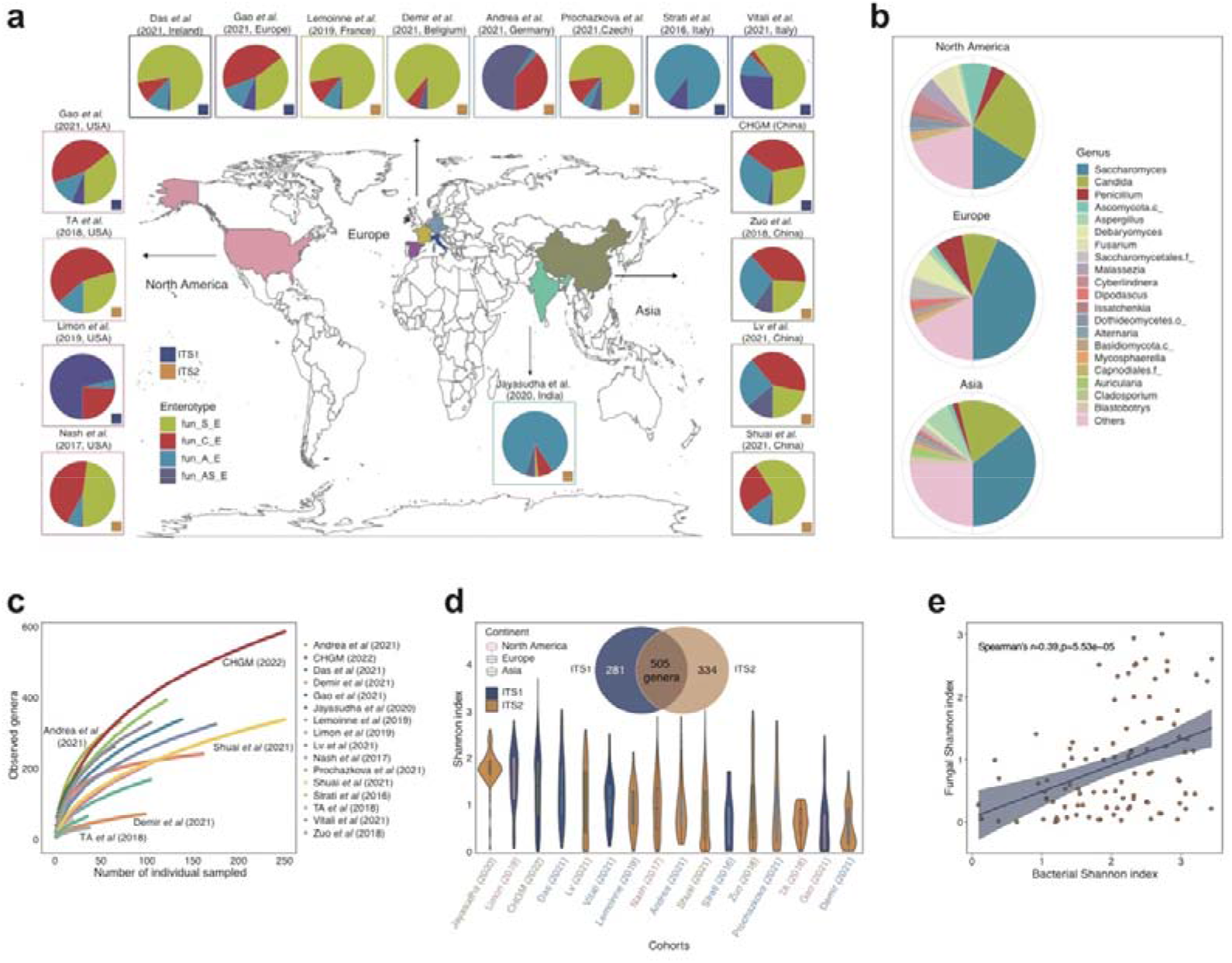
Composition and diversity of the human gut mycobiome across studies and geographic sites. **a**, Geographic distribution of study populations and associated fungal enterotypes, where the datasets are sequenced with either ITS1 or ITS2 barcodes. **b**, Genus-level gut mycobiome composition across the three continents (North America, Europe, and Asia). **c**, Cumulative curves of the number of detected genera according to the number of sequenced samples from different study populations. **d**, The distribution of Shannon diversity across study populations. The Venn diagram shows the number of fungal genera detected by ITS1- and ITS2- based amplification. **e**, The correlation between the Shannon index of bacteria and that of fungi in the Zuo *et al*(*19*) cohort, with shaded region representing 95% confidence intervals of the linear regression.

The gut mycobiome composition and the fungal diversity varied significantly across cohorts, which may be partially attributed to biological and technical factors such as geography and sequencing methods (Fig. 1b-d; *p* < 0.001, PERMANOVA, see Supplementary Note). Though we obtained a total of 1,120 genus-level taxonomic groups after combining all samples, the observed number of the fungal genera was still considerably below the estimated saturation level (Extended Data Fig. 1c), suggesting that a requirement for further increase in sample size to characterize the comprehensive gut fungal diversity. At the genus level, *Saccharomyces* and *Candida* were the most abundant genera across all samples, followed by *Penicillium* and *Aspergillus* (Fig. 1b). These genera are also the most common commensal fungi in other human body sites, including skin, lung, and oral cavity(*29, 30*), indicating their possible well-balanced symbiotic relationship with humans.

The gut mycobiome, compared with the paired bacteriome, demonstrated a significantly lower Shannon diversity yet higher between-individual dissimilarity (Extended Data Fig. 1e). Such observation was in line with the previous studies showing that, in comparison with the gut bacteriome, the gut mycobiome was less diverse but more individual-specific(*21, 31*). In addition, we found a positive correlation between the pairwise dissimilarities of fungal and bacterial communities across studies that had matched mycobiome and bacteriome datasets (Extended Data Fig. 1f), as well as a significant positive correlation between the alpha-diversity indices of the two communities (Fig. 1e; Supplementary Table 3), suggesting the possible between-kingdom interactions of gut microbiota.

### Enterotypes of the human gut mycobiome

To investigate the overall structural and compositional patterns of the human gut mycobiome, we stratified the genus-level fungal compositions of the 3,363 samples into distinct groups, i.e., enterotypes (Methods). The clustering analysis revealed that both ITS1- and ITS2-combined datasets formed four distinct clusters (Fig. 2a, Extended Data Fig. 2a), and these enterotypes were highly concordant across clustering results obtained at other taxonomic levels (Extended Data Fig. 2d). This finding remained unchanged even at a removal of the half samples (Extended Data Fig. 2b). Three of these fungal enterotypes were found in both ITS1- and ITS2-sequencing datasets, where *Saccharomyces, Candida*, and *Aspergillus* were the most abundant genera, respectively (Extended Data Fig. 2e). Therefore, we defined the *Saccharomyces-*dominated enterotype as fun_S_E, and the *Candida*- and *Aspergillus*-dominated enterotypes as fun_C_E and fun_A_E, respectively. In addition to these three enterotypes, we also observed a fourth enterotype in both ITS1 and ITS2 (Fig. 2a). However, the fourth enterotype in ITS1 was dominated by an unclassified *Ascomycota* phylum (*Ascomycota*.*sp*, presented in 15.1% of ITS1 samples), while in ITS2 it was driven by an unclassified *Saccharomycetales* order (*Saccharomycetales*.sp, presented in 5.5% of ITS2 samples). Such a difference observed for the fourth enterotype between ITS1 and ITS2 can be attributed to different amplicon-targeted regions by ITS1 and ITS2. Hierarchical clustering on the combined datasets (ITS1 and ITS2) shows that these two enterotypes can be grouped together, suggesting that these two enterotypes had similar structures (Extended Data Fig. 2c). Thus we defined the fourth enterotype as fun_AS_E hereinafter.

**Fig. 2.**
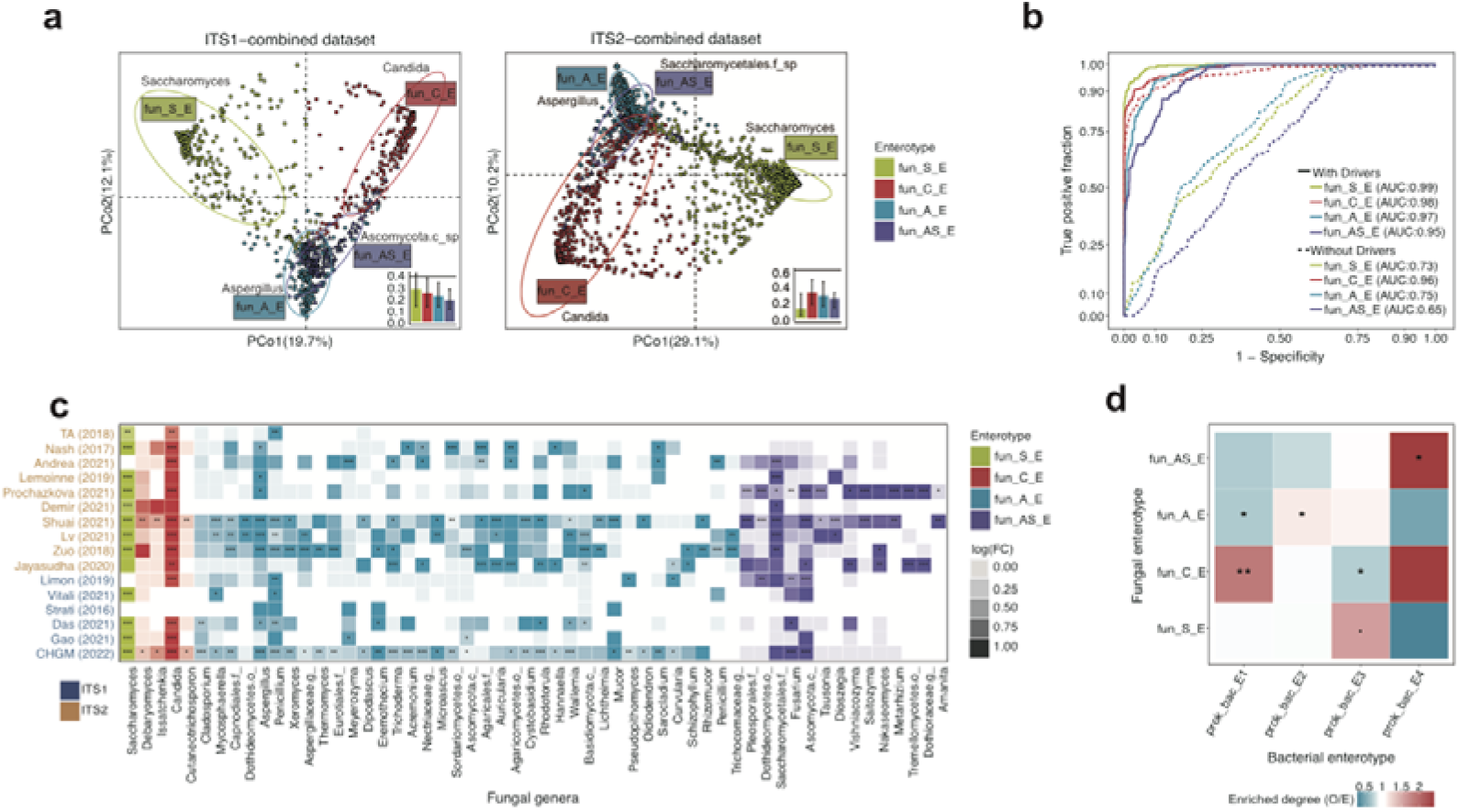
The enterotypes of the human gut mycobiome. **a**, Clustering results of fungal enterotypes on ITS1 and ITS2 datasets and visualized by principal coordinate analysis (PCoA). The between-sample distances within each cluster compared to the median distance between clusters (black line) are shown at the bottom right of each panel. The bar height is the median distance, and the whiskers represent the 25^th^ and 75^th^ quantiles. **b**, A four-enterotype classifier trained on the ITS2-sequencing datasets was applied to predict enterotypes in the ITS1-sequencing datasets. “Without drivers” refers to excluding the driver genera *Candida, Saccharomyces, Aspergillus, Saccharomycetales*.*sp*, and *Ascomycota*.*sp* when training the classifiers. **c**, The concordance of enterotype-associated fungal genera and enrichment trends across different cohorts, and log(FC) denotes the log-transformed fold change of the average relative abundance of the genera within respective enterotypes relative to that of others. Asterisks represent the statistical significance of the multiple testing corrected on-sided non-parametric Wilcoxon test (*adjusted *p* < 0.05, **adjusted *p* < 0.01, ***adjusted *p* < 0.001). **d**, The correlations between fungal enterotypes and bacterial enterotypes in the CHGM cohort. The color reflects the O/E ratio (the ratio of observed count to expected count), and asterisks represent the statistical significance of Fisher’s exact test for each pair of comparison: \**p* < 0.05, \*\**p* < 0.01.

We further confirmed the robustness of the enterotypes by performing a cross-dataset validation analysis between the ITS1- and ITS2-combined datasets with a LASSO logistic regression model (Methods). In the first instance, the model’s high prediction accuracy (Fig. 2b, Extended Data Fig. 3) supported the fungal enterotypes’ robustness. We also obtained a good performance of cross-validation in the absence of these enterotypes’ driver genera, revealing the enterotypes’ ability to characterize the overall fungal community structure independent of the main driver genera (Fig. 2b, Extended Data Fig. 3). Moreover, the consistent enterotype-specific fungal genera profiles across cohorts provided further solid evidence for the robustness of fungal enterotypes (Fig. 2c).

We then examined the geographical and ecological characterizations of the fungal enterotypes. Among the different populations, we found that the fun_C_E enterotype was less common in the European populations (Fisher’s exact test, ITS1: *p* = 4.67e-14; ITS2: *p* = 3.92e-09), while the fun_S_E enterotype was relatively rare in the populations from North America (Fisher’s exact test, ITS1: *p* < 2.2e-16; ITS2: *p* < 2.2e-16). This difference might be partially attributed to the significantly decreased abundance of *Candida* in European populations and that of Saccharomyces in North American populations (Extended Data Fig. 1a). Furthermore, we observed that both the fun_S_E and fun_C_E had the lowest diversity (Extended Data Fig. 2f), and a strong and inverse correlation between the fungal alpha diversity indices and abundances of their respective driver genera (Pearson’s *r* < -0.3, *p* < 2.2e-16).

In addition, we explored the relationship between the fungal and bacterial enterotypes with paired ITS1 for fungal profiling and metagenomics data for bacterial profiling as both data types were available for the CHGM cohort (see methods). Four bacterial enterotypes, which were identified following the same procedure as that of the fungal enterotypes with genus-level metagenomics data (Extended Data Fig. 4), were respectively dominated by *Bacteroides* (20.2% and 37.4% abundances in two bacterial enterotypes, annotated as prok_bac_E1 and prok_bac_E2, respectively), *Prevotella* (42.5% abundance in the prok_bac_E3 enterotype) and *Enterobacteriaceae* (34.9% abundance in the prok_bac_E4). Such observations were in line with those previously reported in the Asian populations(*15, 32*). In addition, we observed significant correlation between the fungal and bacterial enterotypes (Fig. 2d). For example, the fun_C_E fungal enterotype was enriched in the prok_bac_E1 enterotype (*p* = 3.6e-03, Fisher’s exact test) and depleted in the prok_bac_E3 enterotype (*p* = 0.024). We also observed that the fun_A_E enterotype showed a trend to be enriched in the bacterial enterotypes prok_bac_E2, while the fun_AS_E enterotype was enriched in the prok_bac_E4 (both *p* = 0.05). Together with the consistent results from other studies (Extended Data Fig. 5a), such evidence suggested a significant correlation between fungal and bacterial communities.

### Age has a large effect on fungal enterotypes

We then explored the associations between the fungal enterotypes and the hosts’ basic characteristics, including age, gender and BMI. We noticed that age could significantly explain the inter-individual variation of the human gut mycobiome or strongly affected the fungal enterotypes in four cohorts with available age metadata including the CHGM cohort, Gao *et al*(*20*), Limon *et al*(*12*), and Zuo *et al*(*19*) (Fig. 3a, Supplementary Table 4). The insignificant explanatory power of age on the fungal enterotypes in the study by Gao *et al*(*20*) was likely attributable to the small sample size (n=31). As shown in Fig. 3a, fun_C_E (38.8%) and fun_AS_E (34.0%) were significantly enriched in the elderly participants (age > 60 years), while fun_S_E (37.3%) and fun_A_E (44.9%) were significantly enriched in the young participants (age < 30 years, *p* < 0.05, Fisher’s exact test). In addition, a significant inverse correlation between the fungal Shannon diversity and chronological age was observed (Pearson’s *r* = -0.19, *p* = 3.34e-08). Moreover, a multi-variable linear regression analysis on 531 healthy participants from these four cohorts identified 21 fungal genera that significantly correlated with age (Fig. 3b; Methods). Notably, nine age-associated fungal genera were observed to have a different abundance distribution among the three fungal enterotypes (Supplementary Table 5). Among these genera, *Candida*, one driver genera of the fun_C_E, had a positive correlation with age, while two other genera, *Saccharomyces* and *Aspergillus*, showed an inverse trend. This observation was consistent with the age distribution trends of their respective fungal enterotypes (Fig. 3a). Hence, we suspected that the association of fungal enterotypes with age is at least partially driven by their respective dominant fungal genera. No significant association of fungal enterotypes with BMI or gender was found in any cohort (Supplementary Table 4).

**Fig. 3.**
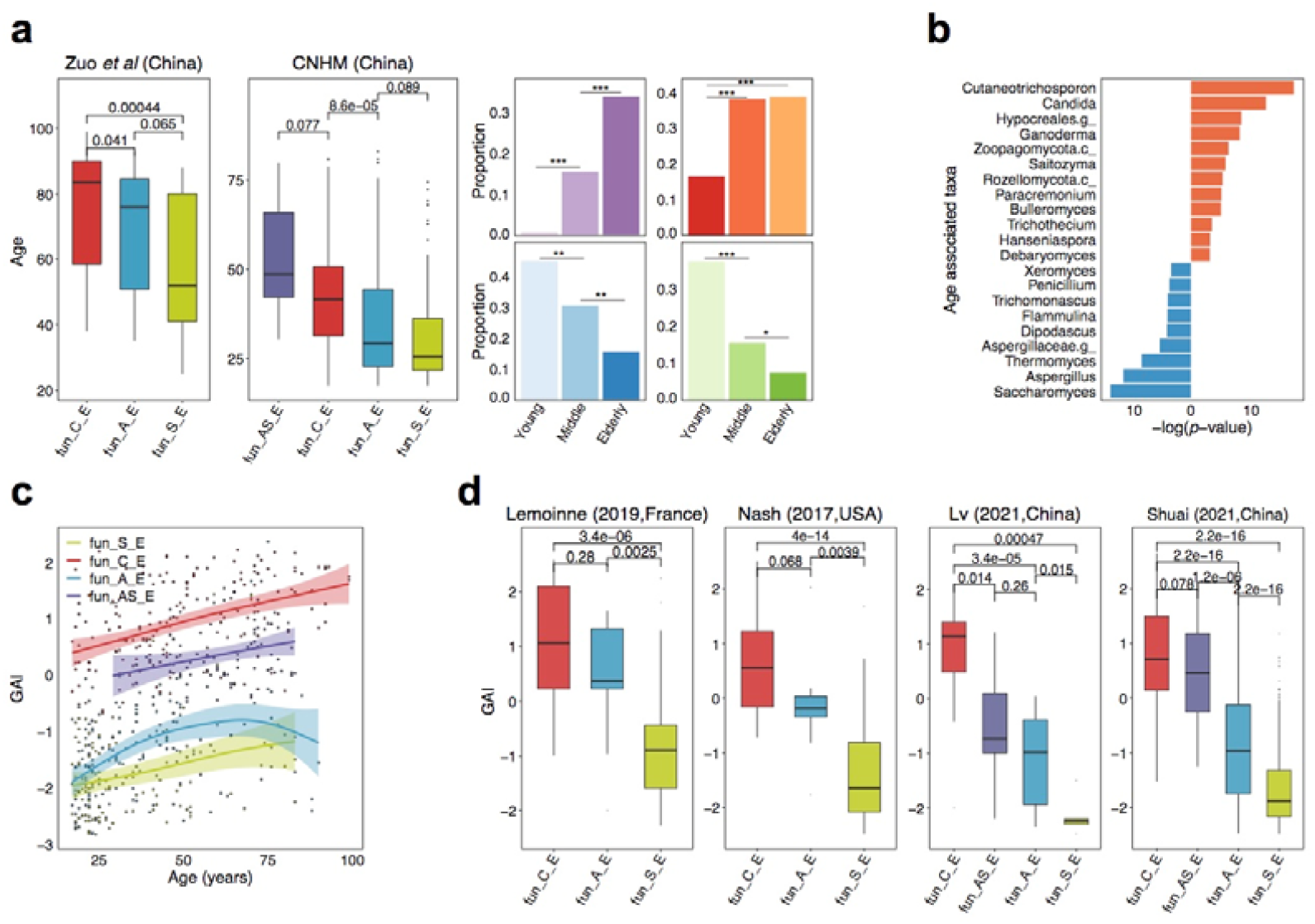
Age distribution and the gut aging indices of fungal enterotypes. **a**, Age distribution of fungal enterotypes in two cohorts from China with *p* values from two-tailed Wilcoxon test *p* values shown for the age difference between enterotypes (left two panels). The right panel shows the proportion of fungal enterotypes in young (18-30 years), middle (31-60 years), and old (>60 years) age groups from these two cohorts, respectively, with asterisks showing the statistical significance of Fisher’s exact test (\**p* < 0.05, \*\**p* < 0.01, \*\*\**p* < 0.001). **b**, The age-associated fungal genera with *p* values < 0.05, where the red bar represents a positive correlation while the blue one represents a negative one. **c**, The correlation between the gut aging index (GAI) and age after the LOESS smoothing for each fungal enterotype on four cohorts with available age data (CHGM cohort, Gao *et al*(*20*), Limon *et al*(*12*), and Zuo *et al*(*19*)). fun_S_E: Pearson’s *r* = 0.30, *p* = 2.1e-03; fun_C_E: Pearson’s *r* = 0.45, *p* = 8.4e-10; fun_A_E: Pearson’s *r* = 0.36, *p* < 3.0e-06; fun_AS_E: Pearson’s *r* = 0.27, *p* = 1.3e-02. **d**, The distribution of GAI across fungal enterotypes in different cohorts. Two-tailed Wilcoxon test *p* values are displayed above the boxplots.

To further quantify the association between the fungal enterotypes and age in other cohorts without available age metadata, we calculated a gut aging index (GAI) for each sample based on the 21 age-associated fungal genera, where higher GAI scores indicating a higher level of intestinal aging (Methods). According to our results, the GAI showed a strong correlation with the age of participants in each enterotypes (Fig. 3c). Of note, participants of the fun_C_E and fun_AS_E enterotypes had consistently higher GAI scores across their lifespan, while those of the fun_S_E and fun_A_E had relatively lower GAI scores (Fig. 3c). Similar results found in healthy subjects of other cohorts without available age metadata further validated the significant associations of GAI scores with the fungal enterotypes (Fig. 3d). Consequently, participants of the fun_C_E enterotypes that contained more age-positively related fungal taxa represented a higher intestinal aging degree, while the physiological condition of the fun_S_E enterotype exhibited a younger state (Fig. 3c,d). Additionally, the distribution of GAI scores in participants with different bacterial enterotypes became another piece of evidence to support correlations between fungal and bacterial enterotypes. For example, participants of the E3_bac enterotype (enriched in fun_S_E) had the lowest GAI scores similar to those of the fun_S_E (Extended Data Fig. 6d). Furthermore, higher GAI scores, as what we observed in patients with intestinal dysbiosis compared to their paired controls, might indicate an occurrence of aging-related pathological changes in the intestine (Extended Data Fig. 6e, Supplementary Note).

### Functional variations across fungal enterotypes

To characterize the bioactive potential of the fungal enterotypes, we annotated fungi-contributed pathways based on the paired shotgun metagenomics data in the CHGM cohort (Methods). In total, we identified 388 biological pathways in the cohort, among which 48 were contributed by fungi alone and 104 were contributed by both bacteria and fungi (fungi-contributed pathways hereafter). Functional richness (the observed number of fungi-contributed pathways) did not vary among fungal or bacterial enterotypes (Extended Data Fig. 2g). However, we identified a total of 31 fungi-contributed pathways whose distribution varied across enterotypes (adjusted *p* < 0.05, Supplementary Table 6). Furthermore, the relative abundances of these pathways were also significantly correlated with those of 14 fungal genera (Fig. 4a, adjusted *p* < 0.05, Supplementary Table 6). An overrepresentation of pathways related to carbohydrate degradation in the fun_AS_E enterotype was observed, suggesting a possible increase in saccharolytic and proteolytic potential (Fig. 4a). Notably, most of the fun_S_E enriched pathways were positively associated with the relative abundance of *Saccharomyces*, which implied the essential roles of genus *Saccharomyces* in these biological pathways. Two pathways involved in heme biosynthesis (PWY-5920 and HEME-BIOSYNTHESIS-II) were enriched in the fun_C_E enterotype and associated with the fun_C_E dominate genera, i.e., *Candida*. It has been demonstrated that heme, the key iron source for pathogenic bacteria, could have a negative impact on the intestinal mucosa and result in a higher risk of colorectal cancer (CRC)(*33, 34*). Thus the participants of fun_C_E enterotype might have an increased risk of developing CRC.

**Fig. 4.**
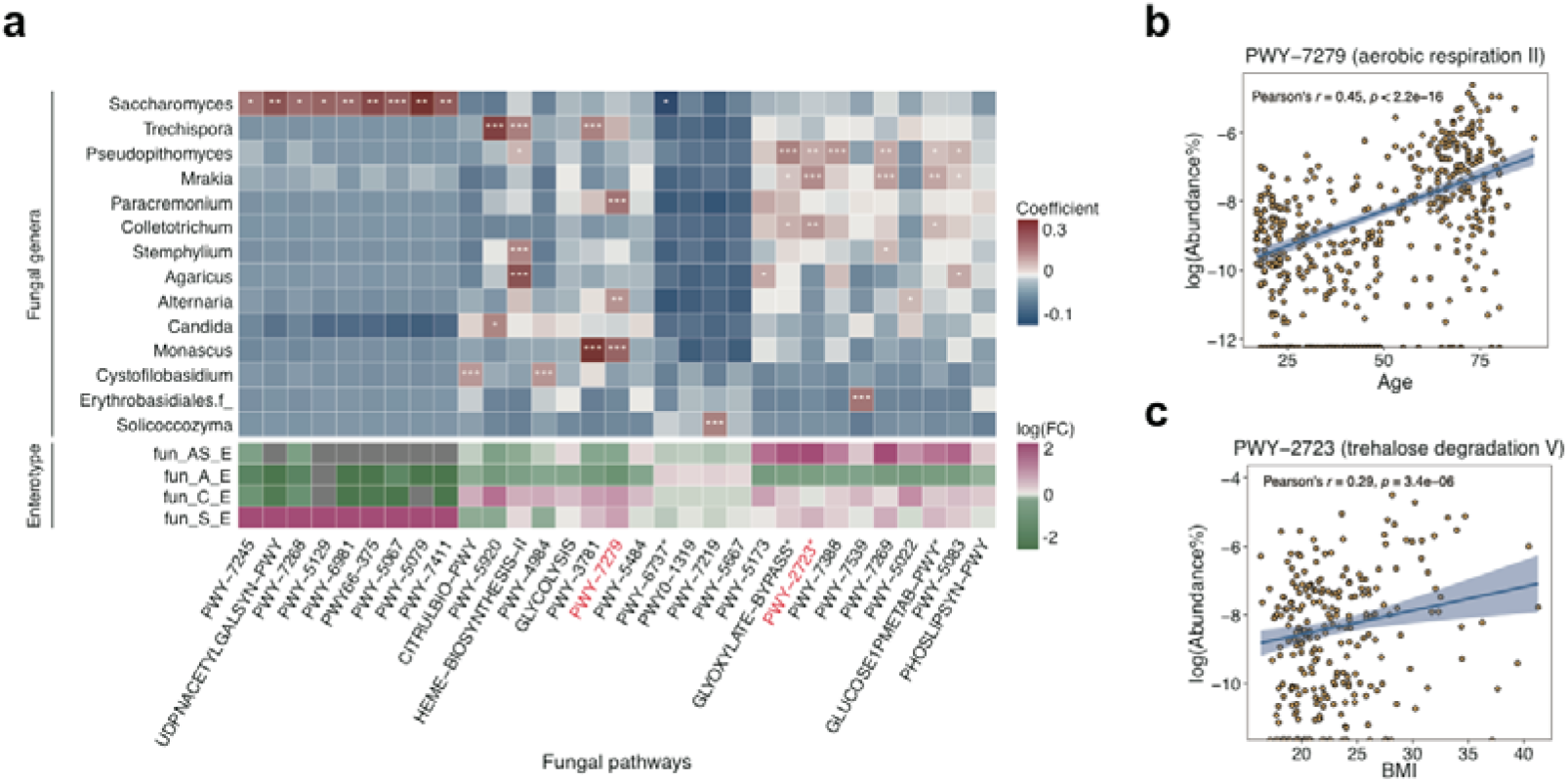
Metabolic pathways associated with fungal enterotypes. **a**, The fungal pathways enriched in different fungal enterotypes (bottom) and associated fungal genera (top), and log(FC) denotes log-transformed fold change of the average relative abundance of the pathway within respective fungal enterotypes relative to that of the others. Asterisks denote multiple testing corrected Pearson correlation tests: *adjusted *p* < 0.05, **adjusted *p* < 0.01, ***adjusted *p* < 0.001. Stars mark the metabolic pathways involved in carbohydrate degradation. **b**, The relationship between the fungi-contributed pathway PWY-7279 and age. **c**, The relationship between the fungi-contributed pathway PWY-2723 and BMI.

To further examine the impacts of fungal enterotypes on human health, we explored these enterotype-associated pathways’ correlations with their host properties. We observed a significant positive correlation between the relative abundance of the fun_C_E-associated pathway PWY-7279 (aerobic respiration) and subject age (Fig. 4b), consistent with the previous observation that the elderly population contained a higher abundance of pathways involved in microbial respiration(*35, 36*). One possible explanation is the higher oxygen level caused by inflammation related to aging promotes aerobic respiration in the gut microbiome(*37*). Additionally, one of the previously detected age-positively related genera, *Paracremonium*, was also shown to be associated with aerobic respiration pathways (Fig. 3b, Fig. 4a). Moreover, we found a significant positive correlation between the host BMI and the PWY-2723, a trehalose degradation pathway (Fig. 4c). The fun_AS_E enterotype, where the PWY-2723 was enriched, had a similar enrichment of biological pathways related to energy metabolism (Fig. 4a). These results are not only consistent with the higher BMI levels in the participants with fun_AS_E enterotype (Extended Data Fig. 6f), but also in line with the previous findings that the microbiota of obese individuals has an increased capacity for energy harvest(*38*). Thus, the functional differences observed across fungal enterotypes can partly explain the host phenotypes variations among fungal enterotypes.

### fun_C_E enterotype is prevalent in disease populations

We further examined the clinical relevance of the fungal enterotypes by assessing their associations with human diseases. By comparing the fungal enterotype’s structure of healthy participants with that of patients with adjustment of age, we found that the fun_C_E enterotype was significantly more prevalent in patients of diseases such as type 2 diabetes, clostridium difficile infection, alcoholic hepatitis, and Alzheimer’s disease (Fig. 5a, *p* < 0.05, odds ratio > 1, Fisher’s exact test). Though there was no significant correlation between fungal enterotypes and other human diseases, we observed similar trends of a higher prevalence of the fun_C_E enterotype in the patients of these diseases (Fig. 5a, odds ratio > 1). In contrast, the other two enterotypes (i.e., the fun_S_E and the fun_A_E) were mainly enriched in the healthy participants (Fig. 5a; odds ratio < 1), except that the fun_S_E was enriched in two viral infectious diseases (H1N1 and COVID-19; Fig. 5a). To further quantify the disease associations across fungal enterotypes, we calculated a Gut Microbiome Health Index (GMHI) as previously described(*39*), and a higher GMHI value indicates a healthier status. Consistent with our expectation, the participants of the fun_C_E enterotype were more likely to have the lowest GMHI value (Fig. 5b), while those of the fun_A_E and fun_S_E enterotypes were more likely to have higher GMHI values. Thus, in addition to its association with higher intestinal aging, the fun_C_E enterotype might also be related to higher disease risk.

**Fig. 5.**
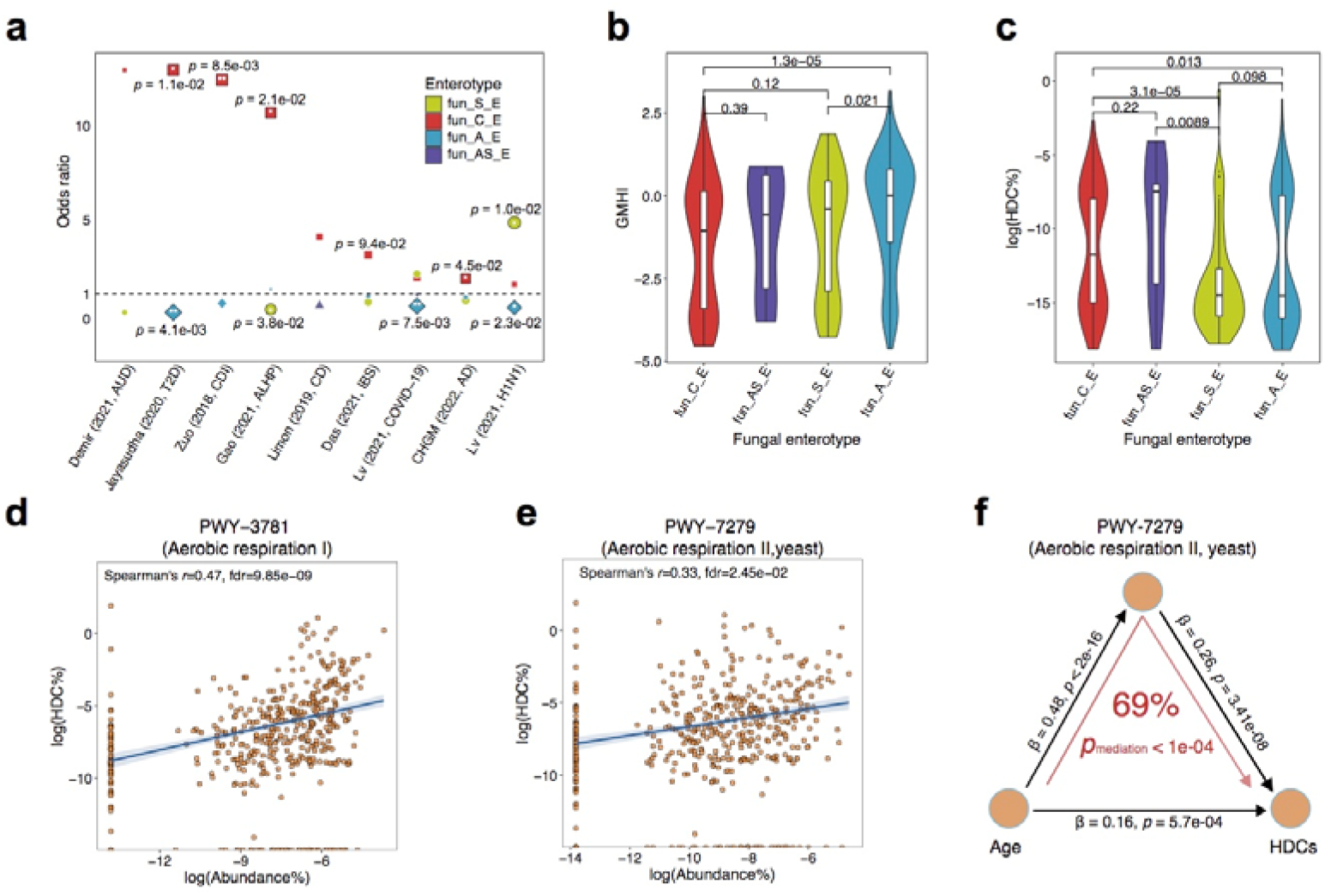
Associations between fungal enterotypes and human diseases. **a**, Enrichment of the fungal enterotypes in human diseases compared to the control group after age was controlled; the odds ratios (OR) and *p*-values of the Fisher’s exact test are shown. AUD: alcohol use disorder; T2D: type 2 diabetes; CDI: clostridium difficile infection; ALHP: alcoholic hepatitis; CD: Crohn’s disease; IBS: irritable bowel syndrome; COVID-19: coronavirus disease 2019; AD: Alzheimer’s disease. **b-c**, Violin plots showing median and quartiles of gut microbiome health index (GMHI) (**b**) and human DNA contents (HDCs) (**c**) across fungal enterotypes in the CHGM cohort, where two-tailed Wilcoxon test *p* values are displayed above the boxplots. **d-e**, Correlations between the HDCs (Y-axis) and the relative abundance of two pathways related to aerobic respiration (X-axis), namely PWY-7279 (**d**) and PWY-7279 (**e**). The shaded region denotes the 95% confidence interval of the linear regression. **f**, Mediation linkages among the chronological age, pathway PWY-7279, and HDCs. *p*_mediation_ was estimated through a bidirectional mediation analysis with 1,000 bootstraps.

To explore the potential molecular mechanism contributing to the association of the fun_C_E enterotype with disease risk, we examined the intestinal barrier function as indicated by human DNA contents (HDCs) in the CHGM cohort (Methods). The HDC acts as an indicator of the compromised intestinal barrier. Previous studies show a significant elevation in HDCs among patients with several intestinal diseases(*40*). We found that the HDCs were significantly higher in the feces of participants of the fun_C_E and the fun_AS_E enterotypes than those of the fun_S_E and the fun_A_E enterotypes (Fig. 5c; *p* < 0.05, Wilcoxon test). This finding was consistent with the GAI scores of these enterotypes (Fig. 3c). Therefore, the compromised intestinal barrier might help to explain the increased disease risk in participants of the fun_C_E. In addition, we also observed significant correlations between the HDCs and the two fungi-contributed pathways involved in aerobic respiration (Fig. 5d,e; adjusted *p* < 0.05). These results strongly indicated significant relationships among the compromised intestinal barrier (hence the increased HDC), gut aging, and the fungal enterotypes’ distribution and bioactive potential. Furthermore, through a bidirectional mediation analysis, we found that the increased age might contribute to the HDC elevation by affecting the abundance of aerobic respiration pathway (69%, *p*_mediation_ < 1e-04; Fig. 5f), which means the increased level of aerobic respiration significantly mediated the relationship between the age and compromised gut barrier.

## Discussion

In this study, we characterized the human gut fungal community structures with a broad spectrum of ITS sequencing samples from 16 cohorts across 11 countries worldwide, including 572 newly ITS-profiled and metagenomically sequenced samples from China. We confirmed the existence of fungal enterotypes that varied in taxonomic and functional compositions and identified four fungal enterotypes, of which the three most common were dominated by *Candida, Saccharomyces*, and *Aspergillus*, respectively, while the fourth appeared more complex with different driver genera in ITS1 and ITS2 analysis, likely due to amplification biases. We noticed that these enterotypes were closely associated with both age and diseases. Particularly, it is noteworthy that the *Candida-*dominated enterotype (fun_C_E) enriched in the elderly population was associated with multiple human diseases accompanied by a compromised intestinal barrier. Additionally, the fun_C_E-associated fungi-contributed aerobic respiration pathway could mediate the association between aging and the compromised intestinal barrier. Thus, our results revealed both the biological and clinical significance of fungal enterotypes and offered a new perspective on host-microbe interactions.

We revealed significant inter-kingdom correlations between gut bacteriome and mycobiome in terms of both community diversity and enterotypes. The *Candida* enterotype (fun_C_E) with the highest disease association displayed a reduced abundance of *Prevotella copri* (Extended Data Fig. 5b), consistent with the previous finding that a lower abundance of *P. copri* in the gut microbiome might indicate intestinal inflammation(*16*). Additionally, one of the Candida species, *C. albicans*, was overrepresented in the fun_C_E, which might result in intestinal dysbiosis and trigger host inflammation(*19*). Previous study demonstrated that commensal anaerobic bacteria, particularly *Firmicutes* and *Bacteroides*, are critical for maintaining *C. albicans* colonization through the activation of two mucosal immune effectors (H1F-1α and LL-37)(*41*). Given the bidirectional interaction between the fungi and bacteria as well as their symbiotic relationship with the human host, a more refined population stratification for both fungal and bacterial enterotypes might be more effective for disease diagnosis. For instance, although no correlation was observed between AD and bacterial enterotypes within the CHGM cohort (*p* = 0.16, Fisher’s exact test, Extended Data Fig. 6h), we observed a lower fungal diversity and a higher prevalence of the fun_C_E enterotype in AD patients (Extended Data Fig. 6g).

We observed significant associations among age, fungal enterotypes, and disease risk. The fungal diversity decreased with increasing age, a similar trend observed for the gut prokaryotic microbiome as reported in previous studies(*36, 42*). A lower diversity of the human gut microbiome is generally indicative of intestinal dysbiosis(*39*), and a gut ecosystem with high species diversity might be more resistant to external environmental interference(*43*). Consistent with these findings, the diversity of the human gut mycobiome was significantly higher in healthy groups than in non-healthy participants (Extended Data Fig. 6g), and the fun_C_E enterotype with lower diversity had a higher disease risk. Therefore, the fungal diversity decreasing with age might suggest a progressive loss of homeostasis in the gut ecosystem. The GAI scores, defined based on age-associated fungal genera, increased in non-healthy participants, implying these fungal genera’s possible involvement in pathogenesis (Extended Data Fig. 6e). We also noticed a correlation between the Eastern Cooperative Oncology Group (ECOG) scores and GAI scores within the CHGM cohort (Extended Data Fig. 6c; Pearson’s *r* = 0.17, *p* = 0.04). These findings supported the previous conclusion on the overlap between aging-related and disease-related deterioration in the gut microbiome(*44*). Therefore, the shared mycobiome alterations might be partly attributable to aging-associated disorders such as frailty and cognitive decline. In addition to the aging-associated pathological changes, the dietary habits, lifestyle, and administration of antibiotics, which can significantly affect our gut microbiome(*45, 46*), also vary during different stages of human life(*47*). Thus, age is associated with a combination of multiple factors, which, in turn, affect fungal enterotypes. Given the occurrence of age-related changes in both the human gut mycobiome and bacteriome, we recommend combining both for future research into the underlying mechanisms of the gut microbiomes during the aging process.

We also noticed several limitations of our study. Firstly, the presence of the fungi detected in the stool samples does not necessarily indicate their long-term colonization in the gut as many of the detected fungi are also commonly found in the food and oral cavities. One longitudinal study of 42 individuals argued that fungi are transient in the human gut and do not colonize in the gut for long periods of time(*48*), but another large-scale study had contrary conclusion and identified several core fungal taxa that were stable over time(*49*). To better unveil the colonization of fungi in the gut, profiling of active fungal community by ITS cDNA analysis is needed in the future work. Secondly, the interactions between the bacteria and fungi were not explored here. The landscape of multi-kingdom interactions can provide insights into the mechanisms underlying the gut mycobiome structure and its association with host physiological conditions. Finally, we explored the functions of gut fungi based on the metagenomics data. However, the metagenomics data is dominated by bacteria, which leads to the underrepresentation of functional profiling of gut mycobiome. Fungi-enriched metagenomics sequencing can be helpful to infer the complete functional profiling of the mycobiome in the future.

## Materials and Methods

### Data collection

We downloaded ITS sequencing data of fecal samples from public databases including National Center for Biotechnology Information (NCBI) sequence read archive (SRA) and China National GeneBank database (CNGBdb). Samples with read number fewer than 10,000 were discarded. Due to the instability and large difference in the human gut mycobiome of infants, we excluded samples from infants. Metadata including demographics (e.g., age, gender, BMI, country) and human disease phenotypes were also retrieved from corresponding publications or databases. As a result, we collected a total of 2,791 public samples from 11 countries covering multiple human disease phenotypes including clostridium difficile infection (CDI), alcohol use disorder (AUD), coronavirus disease 2019 (COVID-19), type 2 diabetes (T2D), irritable bowel syndrome (IBS), alcoholic hepatitis (ALHP), Crohn’s disease (CD) and melanoma. The details for each project including the number of samples, country, associated disease phenotype and used amplicon targets were listed in Supplementary Table 1.

We additionally collected human fecal samples from newly recruited 572 Chinese volunteers (CHGM cohort) with age ranging from 18 to 89 years old, where the fecal mycobiome were profiled with ITS1 amplification. Of these samples, 74 were collected from subjects with Alzheimer’s disease (AD) enrolled in Shanghai Sixth People’s Hospital, whereas others were obtained from healthy volunteers recruited in Wuhan, Shanghai and Zhengzhou. Subjects who take antibiotics, antifungals or probiotics up to 1 month prior to sample collection were excluded from this study. The study protocol was approved by the Human Ethics Committee of the School of Life Science of Fudan University (No, BE1940) and the Ethics Committee of the Tongji Medical College of Huazhong University of Science. All subjects provided informed consent before participation and were asked to complete questionnaires. In total, the combined dataset consisted of 3,363 samples from 16 cohorts and covered 11 countries from three continents, including Europe (615 samples), North America (344 samples) and Asia (2,404 samples); among which, the fungal compositions of six and nine cohorts were determined by ITS1- (960 samples) and ITS2- (2,403 samples) sequencing.

### DNA extraction from fecal samples

After sample collection, the fecal samples from the CHGM cohort were immediately stored on dry ice and transported to a refrigerator at -80°C within 5 hours. Total DNA was extracted from fecal samples using semi-automated DNeasy PowerSoil HTP 96 Kit (Qiagen, 12955-4) according to manufacturer’s instructions. The purified DNAs were quality-checked by 1% agarose gel, and DNA concentration and purity were determined with NanoDrop 2000 UV-vis spectrophotometer (Thermo Scientific, Wilmingtom, USA).

### ITS sequencing and procession

The mycobiome of CHGM cohort was profiled by the sequencing of Internal Transcribed Spacer (ITS), and the ITS1 hypervariable region was amplified with primer pairs ITS1F (5’- CTTGGTCATTTAGAGGAAGTAA-3’) and ITS2R (5’-GCTGCGTTCTTCATCGATGC-3’)(*50*) by an BI GeneAmp® 9700 PCR thermocycler (ABI, CA, USA). The PCR amplification was conducted as follows: initial denaturation at 95°C for 3 mins, followed by 27 cycles of denaturing at 95°C for 30 seconds, annealing at 55°C for 30 seconds, elongation at 72°C for 45 seconds and a final extension at 72°C for 10 mins. The PCR mixtures (20 μL total value) contained 4 μL of 5 × FastPfu buffer, 2 μL of 2.5 mM dNTPs, 0.8 μL of each primer (5 μM concentration), 0.4 μL of FastPfu DNA Polymerase and 10 ng of template DNA. The PCR products were extracted from 2% agarose gel and purified using the AxyPrep DNA Gel Extraction Kit (Axygen Biosciences, Union City, CA, USA) according to manufacturer’s instructions, and further quantified using Quantus™ Fluorometer (Promega, USA). Purified amplicons were pooled and paired-end sequenced on Illumina MiSeq PE300 platform (Illumina, San Diego, USA) according to the standard protocols by Majorbio Bio-Pharm Technology Co. Ltd. (Shanghai, China).

The raw ITS reads were first demultiplexed, quality-filtered by fastp version 0.20.0(*51*) and merged by FLASH version 1.2.7(*52*) with the following criteria: (i) the 300bp reads were truncated at any site with an average quality score < 20 over a 50bp sliding window, and the truncated reads shorter than 50bp were discarded; (ii) only overlapping sequences longer than 10bp were assembled according to their overlapped sequence, and the maximum mismatch ratio of overlap region is 0.2. QIIME2 (version 2019.7) was used for the downstream analysis(*53*). The quality-filtered ITS reads were then denoised and clustered into amplicon sequence variants (ASVs) using DADA2(*54*), and chimeric sequences were identified and removed. Then the Naïve Bayes classifier trained on the UNITE reference database(*55*) was used for taxonomy assignment of individual ASVs. *α*- and *β*-diversity analysis was conducted on samples at the sampling depth of 10,000 by utilizing the R packages “vegan” (version 2.5-7)(*56*) and “phyloseq” (version 1.34.0)(*57*). *α* -diversity was estimated by the Shannon index (evenness and richness of community within a sample), Simpson index (a qualitive measure of community diversity that accounts for both the number and the abundance of features), Faith’s phylogenetic diversity (or Faith’s PD; a qualitative measure of community diversity that incorporates both the phylogenetic relationship and abundance of the observed features) and richness (observed number of features). The fungal genera presented in less than 10 samples were excluded from downstream analysis.

### Metagenomics sequencing and processing

The Fecal bacterial microbiome of CHGM cohort was profiled by whole-genome shotgun sequencing with Illumina HiSeq 2000 platform (Novogen, Beijing, China). DNA libraries were prepared as described previously(*58*). The raw sequencing reads were quality-filtered using fastp version 0.20.0, followed by the use of Bowtie2(*59*) to remove host-derived reads by mapping to the human reference genome (hg38). Quantitative profiling of the taxonomic composition of the microbial communities was performed via MetaPhlAn2(*60*). Profiling of microbial pathways was performed with HUMAnN2 v2.8.1(*61*) by mapping reads to Uniref90(*62*) and MetaCyc(*63*) reference databases. Both the abundance output of MetaPhlAn2 and HUMAnN2 were normalized into the relative abundance. We extracted the metabolic pathways of gut fungi for downstream analysis. The metabolic pathways or bacterial species presented in less than 10 samples were excluded from downstream analysis. To estimate the percentage of human DNA contents (HDCs) within CHGM cohort, we aligned the clean reads to the human reference genome with bowtie2, and the HDCs was calculated as the percentage of mapped reads to the total number of clean reads.

### 16S rRNA sequencing data processing

The 16S rRNA sequencing data available for four cohorts including *Lemoinne* et al(*24*), *Vitali* et al(*64*), *Prochazkova* et al(*27*) and *Zuo* et al(*19*) were downloaded from NCBI SRA. Raw 16S reads were quality filtered, clustered into ASVs and taxonomic annotated using QIIME2 (version 2019.7) as described above. The taxonomies of ASVs were annotated by using the SILVA database(*65*). *α*- and *β*-diversity analysis was conducted on samples at the sampling depth of 25,000. The bacterial genera presented in less than 10 samples were excluded from consideration.

## Ethics approval

This study was approved by the Human Ethics Committee of the School of Life Sciences of Fudan University (No, BE1940) and the Ethics Committee of the Tongji Medical College of Huazhong University of Science and Technology (No, S1241).

## Authors’ contributions

XMZ, YZ, WHC and PB conceived the study and supervised the project. YY, YNP, SCL, JGQ and BHJ managed the sampling and did most of the experiments; SYL, XMZ, YZ, WHC and PB designed the method and performed analysis. S.L wrote the first draft of the manuscript. All authors contributed to the revision of manuscript prior to submission and all authors read and approved the final manuscript.

